# The *Saccharina latissima* microbiome: algal tissue matters more than region, season, and physiology

**DOI:** 10.1101/2022.06.22.497188

**Authors:** Bertille Burgunter-Delamare, Sylvie Rousvoal, Erwan Legeay, Gwenn Tanguy, Stein Fredriksen, Catherine Boyen, Simon M. Dittami

**Affiliations:** CNRS, Sorbonne Université, Integrative Biology of Marine Models (LBI2M), Station Biologique de Roscoff, 29680 Roscoff, France; CNRS, Sorbonne Université, FR2424 Station Biologique de Roscoff, 29680 Roscoff, France; University of Oslo, Department of Biosciences, PO Box 1066, Blindern, N-0316 Oslo, Norway

**Keywords:** holobiont, brown macroalgae, microbiome, metabarcoding

## Abstract

*Saccharina latissima* is a canopy-forming species of brown algae and, as such, is considered an ecosystem engineer. Several populations of this alga are exploited worldwide, and a decrease in the abundance of *S. latissima* at its southern distributional range limits has been observed. Despite its economic and ecological interest, only a few data are available on the composition of microbiota associated with *S. latissima* and its role in algal physiology. We studied the whole bacterial community composition associated with *S. latissima* samples from three locations (Brittany, Helgoland, and Skagerrak) by 16S metabarcoding analyses at different scales: algal blade part, regions, season, and physiologic state.

We have shown that the difference in bacterial composition is driven by factors of decreasing importance: (i) the algal tissues (apex/meristem), (ii) the geographical area, (iii) the seasons, and (iv) the algal host’s condition (healthy vs. symptoms). Overall, *Alphaproteobacteria, Gammaproteobacteria*, and *Bacteroidota* dominated the general bacterial communities. Almost all individuals hosted bacteria of the genus *Granulosicoccus*, accounting for 12% of the total sequences, and eight additional core genera were identified. Our results also highlight a microbial signature characteristic for algae in poor health independent of the disease symptoms. Thus, our study provides a comprehensive overview of the *S. latissima* microbiome, forming a basis for understanding holobiont functioning.

## I. INTRODUCTION

Brown macroalgae, particularly kelps (Laminariales), play essential ecosystem engineering roles in coastal temperate marine environments. Depending on the genus, they are distributed across the western or eastern temperate North Pacific, the Arctic, and North Atlantic Oceans (Bolton, 2010; Araújo et al., 2016). Kelps contribute to primary productivity and are habitat formers providing food and shelter to the local biodiversity (Schiel and Foster, 2006; Schiel and Lilley, 2007). In addition, species of kelps are important in many industries to produce alginates (Peteiro, 2018), human food, medicine (Smit, 2004), or food for abalone aquaculture (McHugh, 2003; Roussel et al., 2019).

*Saccharina latissima* (Linnaeus) C.E. Lane, C. Mayes, Druehl & G.W. Saunders is one of the dominant kelp-forming species of brown macroalgae in Europe. Its tissue growth starts from the meristematic region at the base of the blade, with the older tissue being at the apex part. These older parts can undergo erosion due to senescence and host a higher bacterial diversity, as shown in previous research on other *Laminariales*, notably *Laminaria digitata* (Corre and Prieur, 1990), *Laminaria hyperborea* (Bengtsson et al., 2010), *Laminaria longicruris* (Laycock, 1974), *Laminaria pallida* (Mazure and Field, 1980), and *Laminaria setchellii* (Lemay et al., 2021).

In recent years, a decrease in the abundance of *S. latissima* at its southern range limits has been observed (Araújo et al., 2016; Smale, 2020). The exact processes driving this decline are not fully understood, but it is likely that changes in peak temperature associated with changes in the microbiota might be at least partially linked to this process, as is the case with corals (Bourne et al., 2008; Bosch and Miller, 2016; Peixoto et al., 2017).

Indeed, macroalgal functioning needs to be seen as the result of the interactions between the algal hosts and their associated microbiota, constituting a singular entity termed the algal holobiont (Egan et al., 2013). It has been shown that macroalgal health, fitness, pathogen resistance (Wiese et al., 2009), acclimation to a changing environment (Dittami et al., 2016), and metabolism (Burgunter-Delamare et al., 2020) are regulated and supported by bacterial partners (Goecke et al., 2010). Considering the biofilm composition and deciphering the interactions within the holobiont is thus essential to fully understand the biology of algae. Previous studies were carried out on microbiota of different kelp species like *L. digitata* (Ihua et al., 2020), *L. hyperborea* (Bengtsson et al., 2010), *L. religiosa* (Vairappan et al., 2001) and *L. setchellii* (Lemay et al., 2021), but little is known about the *S. latissima* microbiota. Notably, Staufenberger et al. (2008) analysed the bacterial composition of *Saccharina* from two locations and seasons (Baltic and North Sea; January and April 2006) using denaturing gradient gel electrophoresis (DGGE) and 16S rRNA gene clone libraries. Later, Tourneroche et al. (2020) used 16S metabarcoding and FISH to decipher the bacterial microbiota of young tissues of *S. latissima* sampled in Scotland on one date.

In the present study, we compared the microbiota composition of young *S. latissima* samples from several locations in the Atlantic Ocean (Brittany, Helgoland, and Skagerrak) by 16S metabarcoding analyses to decipher if the microbiota is specific to the area of origin, seasonality, and algal blade part (apex/meristem). We examined the microbiota composition of healthy and diseased algae, aiming to identify possible microbial signatures characteristic of algae in poor health.

## II. MATERIAL & METHODS

### 1. Biological material & Environmental Variables

*S. latissima* were sampled at different sites and dates (**Table 1**). Briefly, samples were taken from three regions (Brittany, Helgoland, and Skagerrak) at low tide (or diving when necessary). Among young individuals (<1m length), five healthy algae and five with physical symptoms (holes, bleaching, twisted blades) were selected for each sampling session. We focused on a general “symptoms” category rather than on a specific disease because it was impossible to find enough individuals with the same symptoms throughout the sampling sessions and sites. The algal material was immediately placed in sterile plastic bags and rapidly (<3h) transported to the laboratory in a cooling box at ca. 4°C.

**Table 1-.** Sampling dates and sites.

| *Places* | *GPS coordinates* | *Dates* | *Time* | *Types of samples* |
| --- | --- | --- | --- | --- |
| Roscoff, Brittany, France | 48°43'47.0 "N 4°00'17.1" W | 10 October 2018<br>23 January 2019<br>18 April 2019<br>31 July 2019<br>29 October 2019 | At low tide mid-day<br>for all | Healthy + Symptoms |
| Helgoland, North Sea,<br>Germany | 54°10'47.3 "N 7°54'59.4" E<br>54°11'27.1 "N 7°52'02.4" E | 10 July 2019<br>11 July 2019 | Diving<br>Low tide | Healthy + Symptoms |
| Skagerrak, Norway | 58°15'15.5 "N 8°31'22.2" E<br>58°22'05.1 "N 8°44'04.7" E<br>58°05'39.3 "N 6°34'54.9" E | 16 October 2018<br>18 October 2018<br>1 April 2019 |  | Healthy samples<br>No samples with<br>symptoms |

Two parts of the blades were sampled: the basal meristem and the tip (**Figure 1**). A disc with Ø2cm was punched out for each part of the blade and placed in a 15 ml Falcon tube containing 5ml of clean silica gel (2-6mm; VWR). Tubes were stored at room temperature for up to 15 days before DNA extraction.

**Figure 1.**
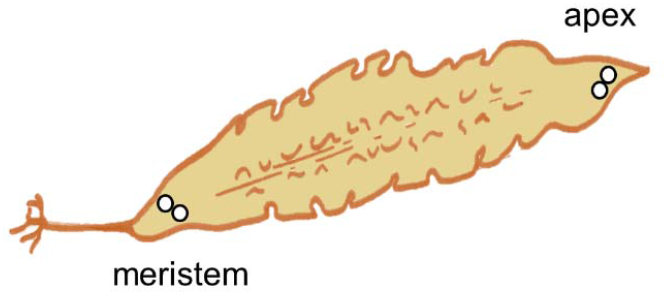
Sampled parts of the *Saccharina latissima* thallus. Two discs (Ø2cm) were punched out in immediate proximity for each part of the blade.

For the samples from Brittany, corresponding environmental variables (temperature, salinity and ammonium, nitrites, nitrates, and phosphate concentrations) were obtained from the Service d’Observation en Milieu Littoral (SOMLIT) database (https://www.somlit.fr/mysomlit/; Astan point; approximately 3.6km North-East of the sampling point). They are available in **Figure S1**.

### 2. DNA extraction

DNA extraction was carried out with the silica-gel stored samples, according to the protocol described by Bernard et al. (2017). Briefly, samples were freeze-dried, and ½ of a disk was ground using a Qiagen TissueLyser II bead beater (3 sessions, 45sec, 30Hz, 3mm stainless steel beads). Nucleic acids were then extracted using a 2% CTAB extraction buffer (100 mM Tris-HCl [pH 7.5], 1.5 M NaCl, 2% CTAB, 50 mM EDTA [pH 8], 50 mM DTT; shaker 250 rpm at room temperature). Supernatants were purified with one volume of chloroform/isoamyl alcohol (24:1) followed by 15min centrifugation at 10 000 rpm (16°C). The upper phase was transferred to a new tube, and ethanol (0.3 vol) was added drop by drop until polysaccharide precipitation was visible, followed by a second chloroform/isoamyl alcohol extraction and recovery of the aqueous phase. The pre-purified DNA was purified using Nucleospin plant II columns (Macherey-Nagel, Germany) according to the manufacturer’s instructions. Finally, DNA was eluted in 50μl of elution buffer (Macherey-Nagel). Blank extractions were also performed, and these extracts were used to identify potential contaminations introduced during the extraction and downstream processing of the samples.

### 3. 16S Metabarcoding

The bacterial community composition associated with algal cultures was determined by 16S metabarcoding. A mock community comprising a mix of DNA from 26 cultivated bacterial strains (Thomas et al., 2019) and negative control were run add treated in parallel to the DNA extracts. For all of these samples, the V3 and V4 regions of the 16S rDNA gene were amplified using the NOCHL primers including Illumina adapters (Thomas et al., 2019), to avoid plastid DNA amplification. Then a standard Illumina protocol for metabarcoding (Illumina, 2013) was run using the Q5^®^ High-Fidelity PCR Kit (New England BioLabs, MA, USA), the AMPure XP for PCR Purification Kit (Beckman Coulter, Brea, CA, USA), and the Nextera XT DNA Library Preparation Kit (Illumina, San Diego, CA, USA). Libraries were quantified with a Quantifluor® ds DNA System (Promega, WI, USA), and mean fragment size was determined using a LabChip® GX Touch™ (Perkin Elmer, MA, USA). An equimolar pool of all samples was generated at a concentration of 4 nM, diluted to 3 pM, spiked with 10% PhiX (Illumina), and sequenced on an Illumina MiSeq sequencer at the Genomer platform (Station Biologique de Roscoff) using a MiSeq v3 kit (2×300bp, paired-end). Raw Illumina reads were deposited at the European Nucleotide Archive under project accession number PRJEB47035.

### 4. Analyses

Sequence analysis was performed using the DADA2 1.14.0 package (Callahan et al., 2016) on R 3.6.2 following the protocol established by Benjamin Callahan (https://benjjneb.github.io/dada2/tutorial.html). Sequences were filtered, allowing for a maximum of 2 expected errors and reducing the read length to 291 bp for forward reads and 265 bp for reverse reads. An amplicon sequence variant (ASV) table was constructed, and chimaeras were removed. The taxonomy of the remaining ASVs was assigned using the Silva_SEED 138 database. The resulting abundance table and taxonomic classification were analysed using Phyloseq 1.30.0 (McMurdie and Holmes, 2013). ASVs that were more abundant in the blank samples than in the algal samples, organellar and eukaryote reads, rare ASVs (<0.01% of total reads), and samples with less than 7688 remaining reads were removed. Non-Metric Multidimensional Scaling analyses (NMDS) were carried out using the Bray-Curtis dissimilarities and the Vegan R package. The most important factor separating the samples in the NMDS was then further explored. The Shannon H diversity index was calculated using Past version 4.02 (Hammer et al., 2001). Statistical analysis of differential abundance was performed at the genus level using ANCOM-BC version 1.4.0 (Lin and Peddada, 2020) with default parameters. Binomial tests followed by a Benjamini and Hochberg (BH) correction (Benjamini and Hochberg, 1995) were carried out to determine the overrepresented genera among the ASVs identified by ANCOM-BC. Then, the factor in question was eliminated from the dataset if possible (grouping of apex and meristem sample, focus on specific region), and the analyses were repeated to determine the next factor. The bacterial core was determined at the genus level and defined as genera present in 90% of replicates for each algal part, season, and location.

## III. RESULTS

### 1. General taxonomy

16S metabarcoding analyses were carried out for all control and algal samples. A total of 4,028,372 raw sequences were generated and, after filtering, assembled into 1,658,746 merged contigs. The taxonomic assignation of mock samples was consistent with the mock composition, and a total of 18,028 ASVs were identified in the dataset. The sequences obtained corresponded predominantly to *Alphaproteobacteria* (34,1% of total reads), followed by *Gammaproteobacteria* (29,5% of total reads) and *Bacteroidota* (26% of total reads).

### 2. Comparison of apex and meristem samples

Global NMDS analysis of all samples demonstrated a clear separation between the apex and meristem samples (**Figure 2A)**. Overall, alpha diversity, as calculated using the Shannon H index (**Figure 2B**), was higher in apex samples than in meristem samples (*p-value* <*0.0001*). Several phyla were found to differ significantly in relative abundance between the apex and meristem samples. The *Actinobacteriota, Firmicutes*, and unclassified *Proteobacteria* (*p*<*0.0001*) were found in higher relative abundance in the meristem samples. The *Alphaproteobacteria* (*p*=*0.00016*), *Bacteroidota* (*p*=*0.00372*), and *Planctomycetota* (*p*=*0.004*) phyla were relatively more abundant in the apex samples (**Figure 2C**). ANCOM-BC analyses revealed a total of 122 ASVs to differ significantly (adjusted *p-value*< 0.05) in relative abundance between the apex and meristem samples (28 ASVs were more abundant in apex and 94 in meristem samples; **Table S1**). The taxonomic groups overrepresented (adjusted *p-value*< 0.05; BH correction) among these significant ASVs are shown in **Table 2:** one genus was significantly overrepresented in the apex samples (*Ki89A_clade*, 39%) and six in the meristem samples (including *Gammaproteobacteria*, 70%; **Table 2)**. The bacterial core in the apex and meristem samples comprises the four genera *Granulosicoccus, Litorimonas, Hellea*, and *Blastopirellula*, accounting for 32% of the total reads for all samples. Five additional genera were systematically present in the apical part: *Algitalea, Arenicella, Portibacter, Tenacibaculum*, and *Bdellovibrio* and accounted for 15% of the total reads.

**Figure 2.**
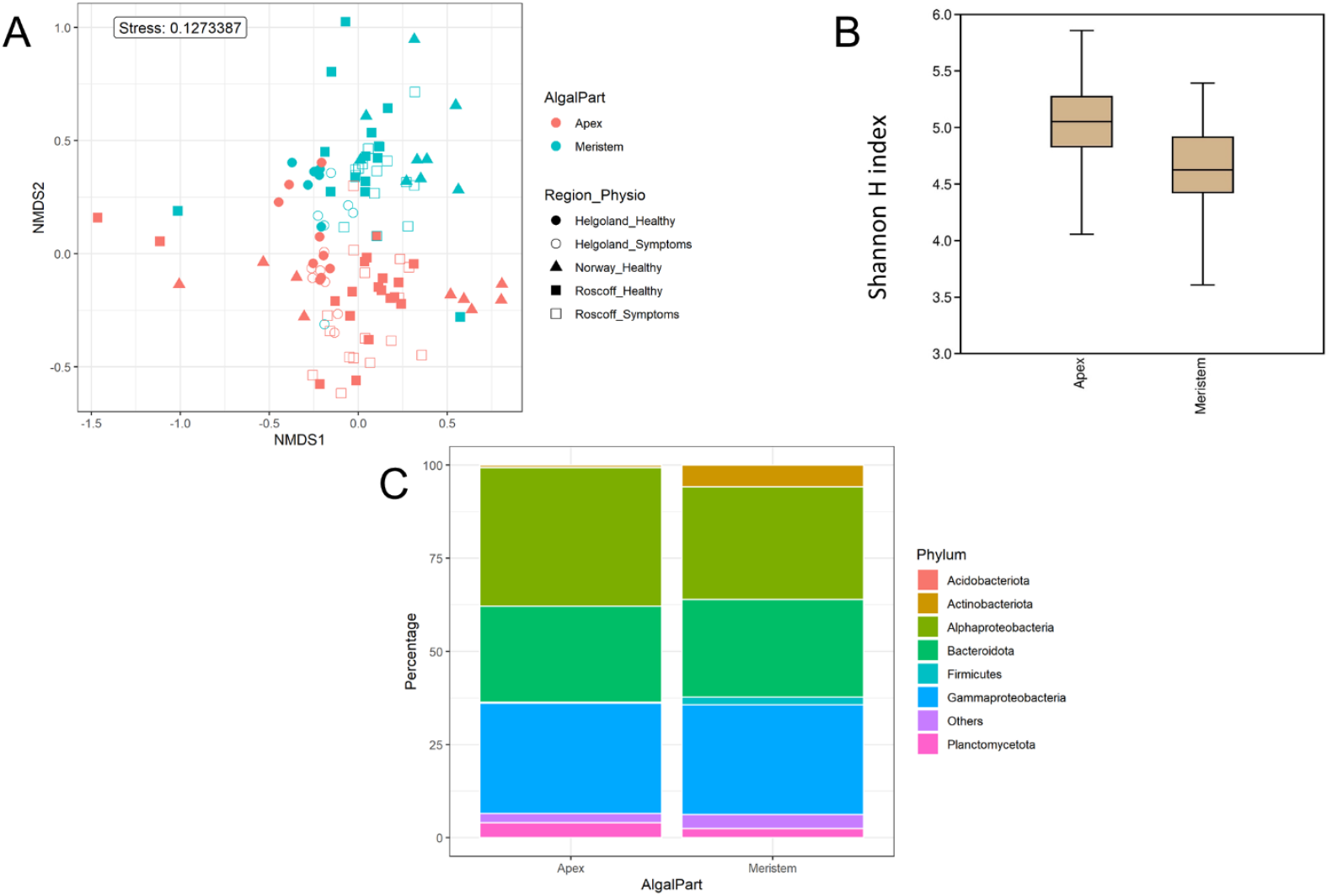
Algal blade part analysis. A) NMDS analysis of the microbiome composition. Results show a clear separation of the apex and meristem samples. B) Box plot of alpha-diversity (Shannon H index) across different sample types. *p-value* <*0.0001*. C) Comparison of microbiome composition between apex and meristem samples. Distribution of 16S rRNA gene metabarcoding sequences per phylum. *Proteobacteria_NA:* not classified as *Alpha*- or *Gammaproteobacteria*.

**Table 2.**
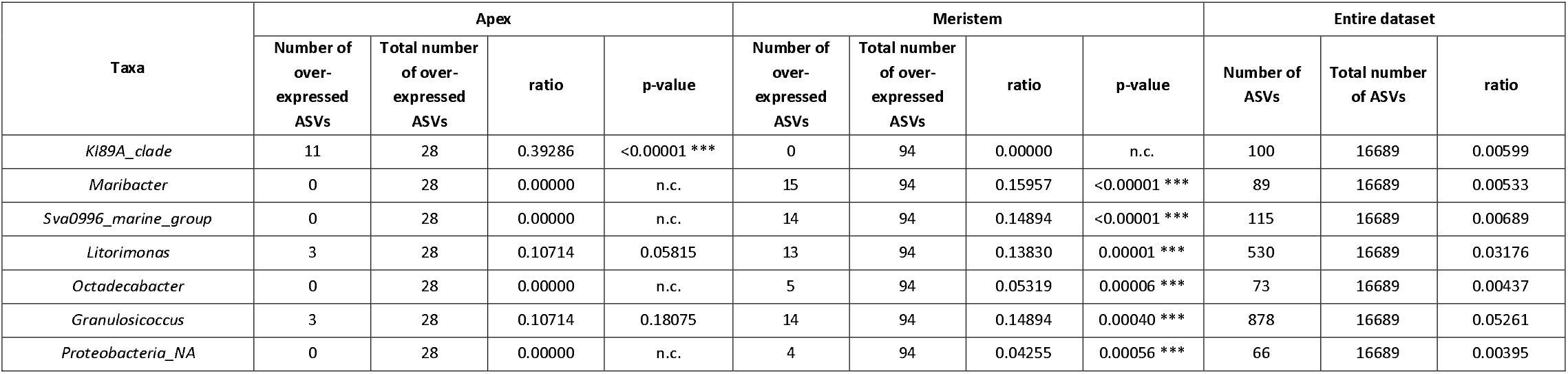
Taxonomic affiliations of the ASVs characteristic for apex and meristem samples, compared with their occurrence in the entire dataset. n.c.: not calculated. P-values are the results of a binomial test, and ***, **, and *indicate that these p-values were significant (*i.e*., the genus is significantly overrepresented among the genera regulated by tissue type) even after a Benjamini and Hochberg correction.

### 3. Comparison of regions

For the following analyses, reads from the apex and meristem samples of the same alga were pooled as individuals to remove the apex/meristem effect. On the NMDS plot, the samples are now grouped according to their region of origin (**Figure 3A**). The alpha diversity did not differ significantly between the regions (**Figure 3B)**. However, at the phylum level, the *Firmicutes* and unclassified *Proteobacteria* were underrepresented in the Norwegian samples compared to Roscoff (p=0.013 and p<0.0001) and Helgoland (p=0.004 and p<0.0001; **Figure 3C)**. *Bacteroidota* and *Alphaproteobacteria* exhibited significantly higher relative abundance in Roscoff than in Helgoland (p= 0.003 for both phyla). At the ASV level, 234 ASVs were represented in higher proportions in the Roscoff samples, 243 in the Helgoland samples, and 18 in the samples from Southern Norway (**Table S1)**. The taxonomic affiliation of significantly over-expressed ASVs (adjusted *p-value*< 0.05; BH correction) is shown in **Table 3**, and six genera were significantly overrepresented in Helgoland samples, one genus in the Norwegian samples (*Rhizobiaceae_NA*), and nine genera in Roscoff samples (*Proteobacteria* 77%; **Table 3)**.

**Figure 3.**
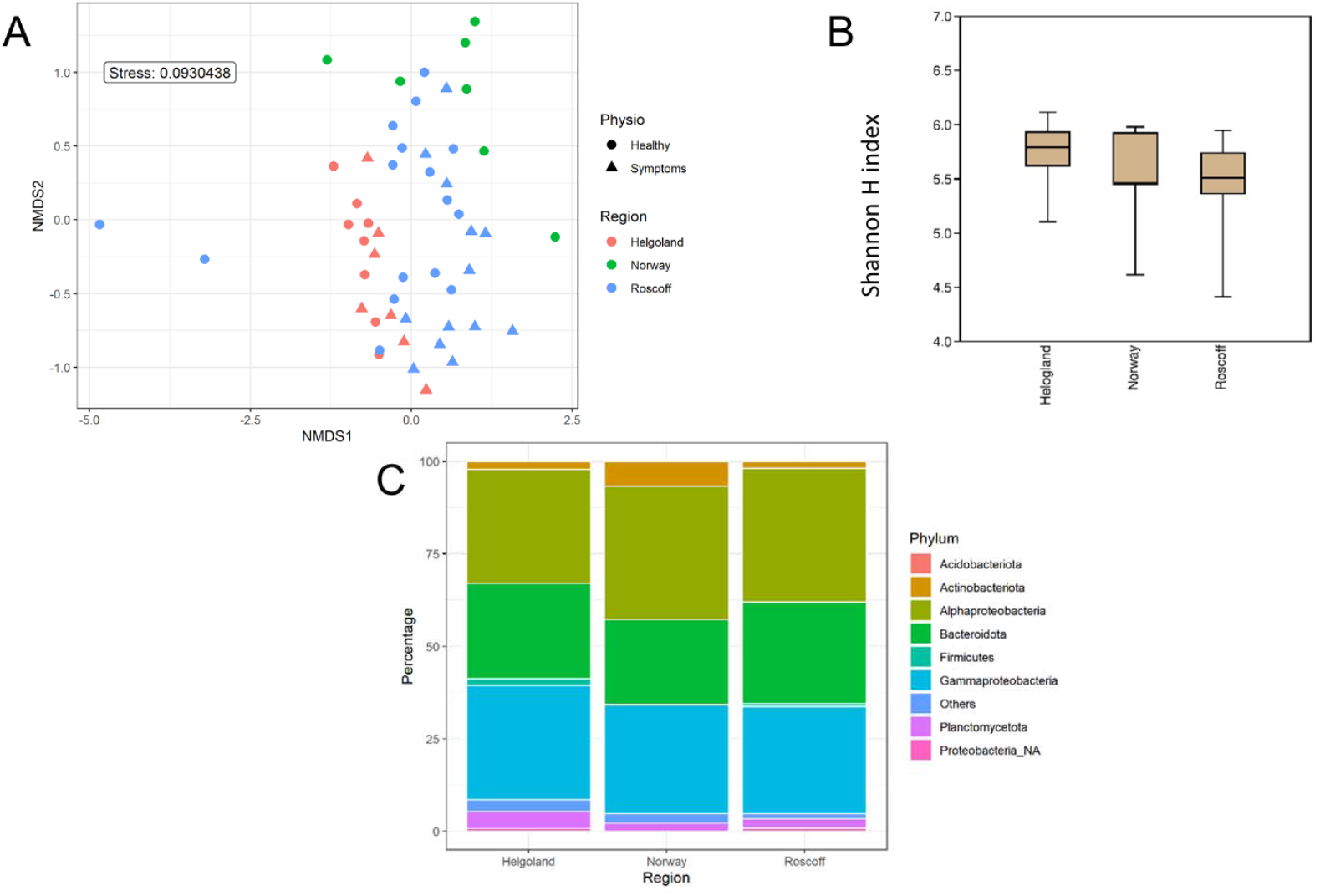
Regions analysis. A) NMDS analysis of the microbiome composition. Results show a clear separation of the samples according to their origin. B) Box plot of alpha-diversity (Shannon H index) across different sample types. *p-value*>0.05. C) Microbiome composition of samples from Helgoland, Norway, and Roscoff. Distribution of 16S rRNA gene metabarcoding sequences per phylum. *Proteobacteria_NA*: not classified as *Alpha*- or *Gammaproteobacteria*.

**Table 3.**
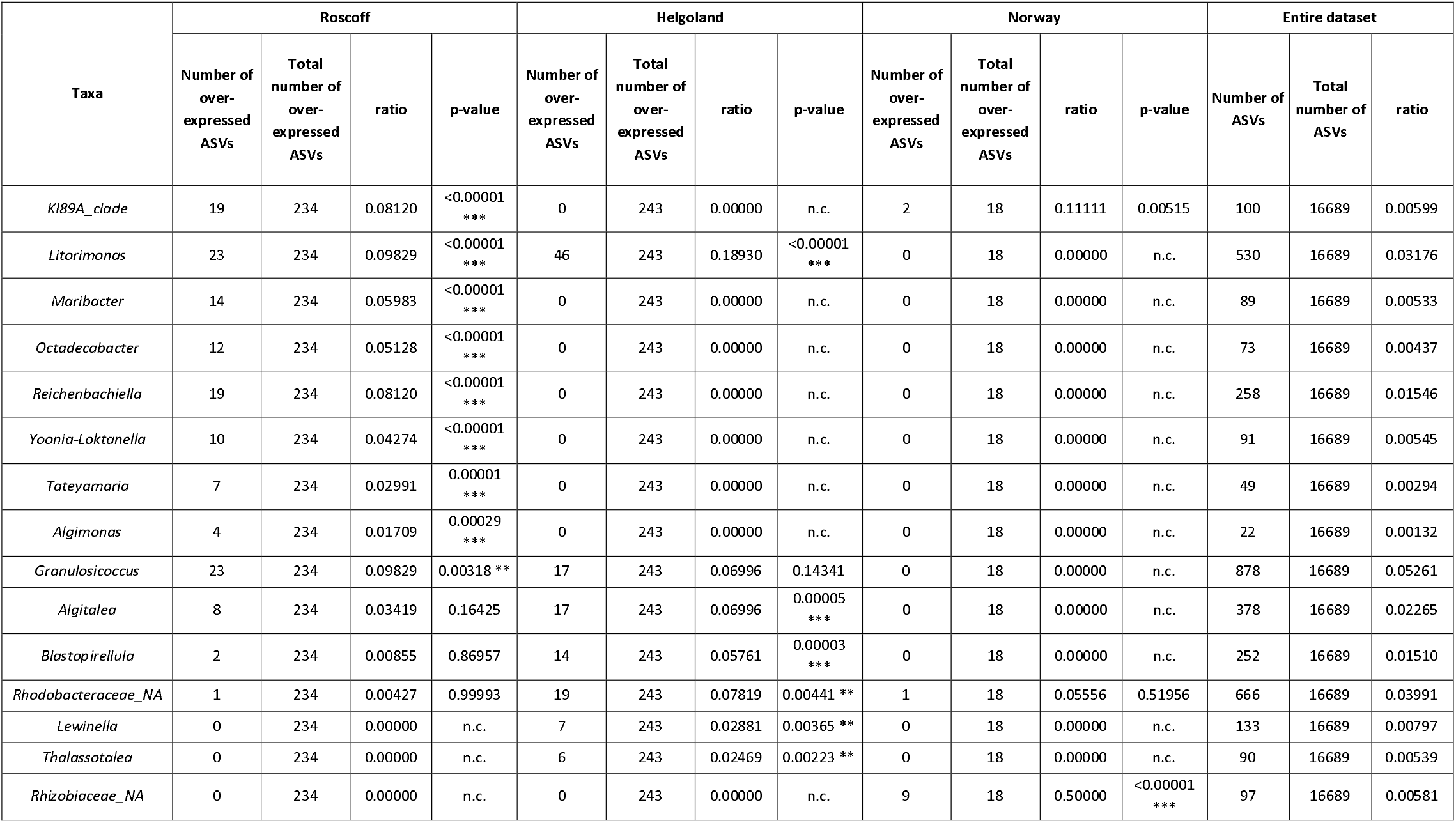
Taxonomic affiliations of the ASVs characteristic for the Roscoff, Helgoland, or Norway samples, compared with their occurrence in the entire dataset. n.c.: not calculated. P-values are the results of a binomial test, and ***, **, and *indicate that these p-values were significant (*i.e*., the genus is significantly overrepresented among the genera regulated by tissue type) even after a Benjamini and Hochberg correction.

### 4. Seasonality

Only Roscoff samples were used to assess the impact of season on the microbiome because these were the only samples with four sampling points from different seasons available. The NMDS analysis shows a separation between the season’s samples. The autumn and winter samples clustered together, and the spring and summer samples were placed on the sides (**Figure 4A),** even if the alpha diversity (**Figure 4B**) did not differ significantly between the seasons. *Actinobacteria* were exclusively found in summer times. *Firmicutes* (p=0.032) were more abundant in autumn samples than in spring samples (p=0.032). *Alphaproteobacteria* were significantly more abundant in autumn than summer (p=0.0017) and winter (p=0.014). *Gammaproteobacteria* were significantly more abundant in spring than in autumn (p=0.032) and winter (p=0.022; **Figure 4C)**. ANCOM-BC analyses revealed 422 ASVs with higher relative abundance in one or several seasons. 126 ASVs were most abundant in winter samples, 85 ASVs in spring samples, 95 ASVs in summer samples, and 115 ASVs in autumn samples (**Table S1)**. The taxa significantly overexpressed among these ASVs (adjusted *p-value*< 0.05; BH correction) are shown in **Table 4**. Most ASVs with higher relative abundance in winter, spring, and autumn samples belonged to the *Alphaproteobacteria* (17%, 24%, and 33% of ASVs). In summer, six genera were over-represented, and 33% of ASVs belonged to the *Gammaproteobacteria*. Also, the Sva0996_marine_group (*Actinobacteria;* 12%) was overrepresented only in these samples.

**Figure 4.**
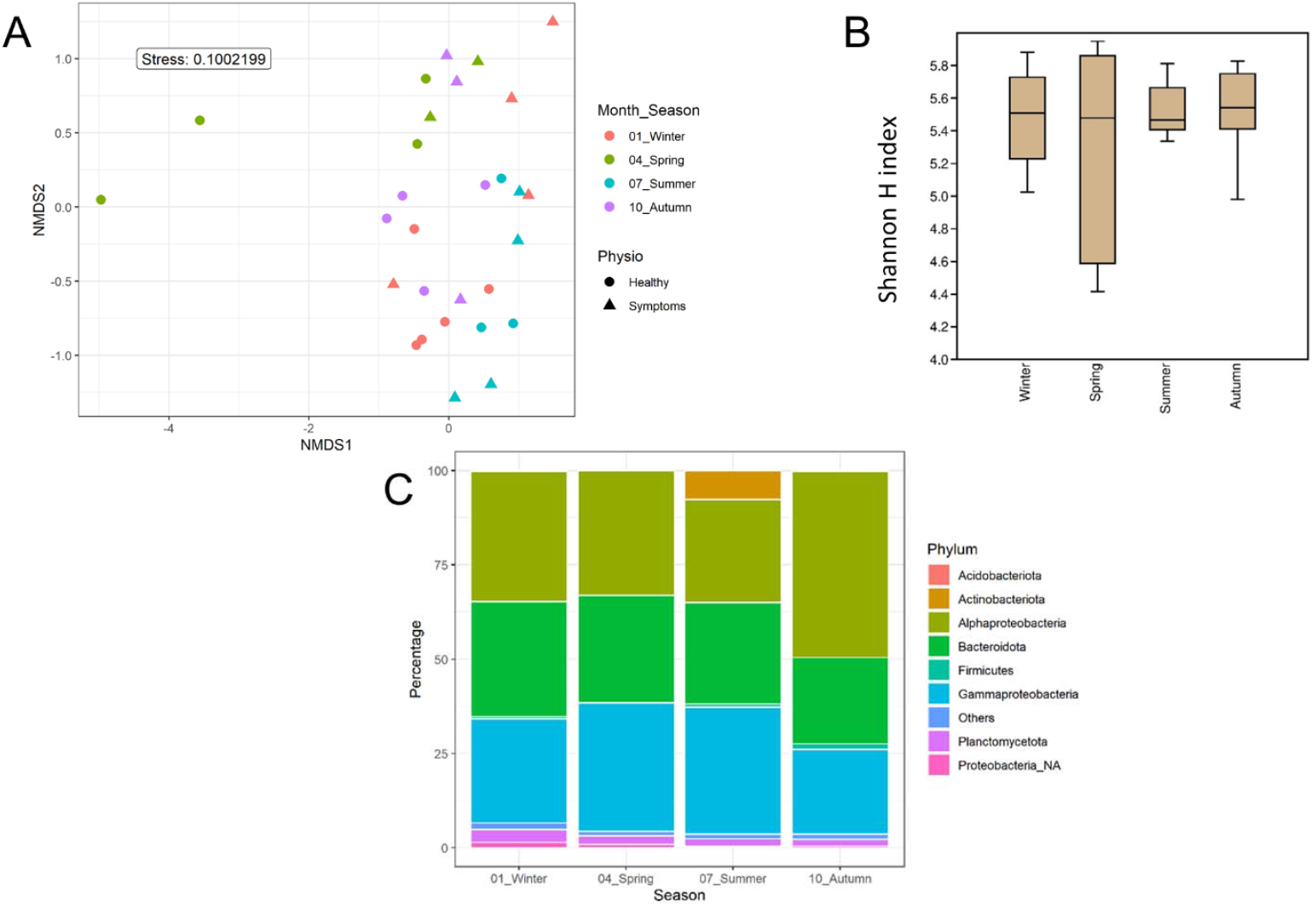
Season analyses. A) NMDS analysis of the microbiome composition. Results show a separation of the samples depending on the sampling’s season. B) Box plot of alpha-diversity (Shannon H index) across different sample types. *p-value*>0.05. C) Seasonal microbiome composition in Roscoff. Distribution of 16S rRNA gene metabarcoding sequences per phylum. *Proteobacteria_NA*: not classified as *Alpha*- or *Gammaproteobacteria*.

**Table 4.**
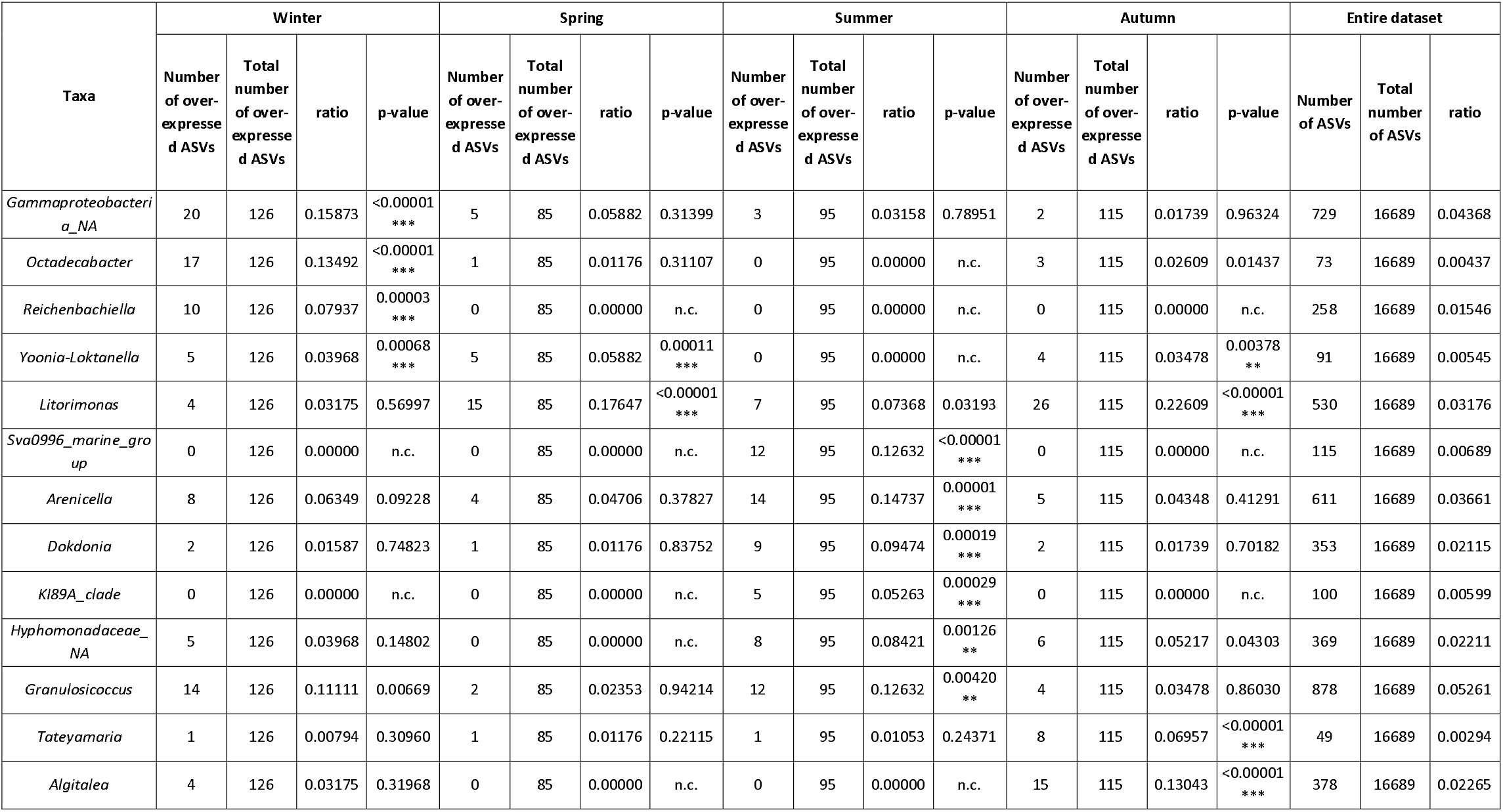
Taxonomic affiliations of the ASVs characteristic for each season from Roscoff samples, compared with their occurrence in the entire dataset. n.c.: not calculated. P-values are the results of a binomial test, and ***, **, and *indicate that these p-values were significant (*i.e*., the genus is significantly overrepresented among the genera regulated by tissue type) even after a Benjamini and Hochberg correction.

### 5. Comparison Healthy / Symptoms

Both healthy samples and samples with symptoms were found only in Roscoff and Helgoland, and the symptoms were diverse: holes, twisted blade, and bubbling in the blade (**Figure 5)**. The NMDS shows no separation between the healthy individuals and those with symptoms (**Figure 6A),** although the Shannon H index indicated slightly higher alpha diversity in algal samples with symptoms than in healthy samples (*p-value=0.046;* **Figure 6B)**. No phyla significantly and systematically differed between healthy algae and algae with symptoms (**Figure 6C)**. This observation also remains true when we distinguish samples between the different types of symptoms (**Figure 6D),** and the samples are still separated depending on the region. However, ANCOM-BC analyses revealed 9 ASVs that were characteristic in either of the groups: four ASVs were more abundant in samples with symptoms (*Alteromonadaceae_NA, Octadecabacter* sp., *Tenacibaculum* sp., *Yoonia-Loktanella* sp.) and five that were more abundant in the healthy ones (*Escherichia/Shigella* sp., *Granulosicoccus* sp., *KI89A_clade, Rhodobacteraceae_NA, Zobellia* sp.; **Table S1**).

**Figure 5.**
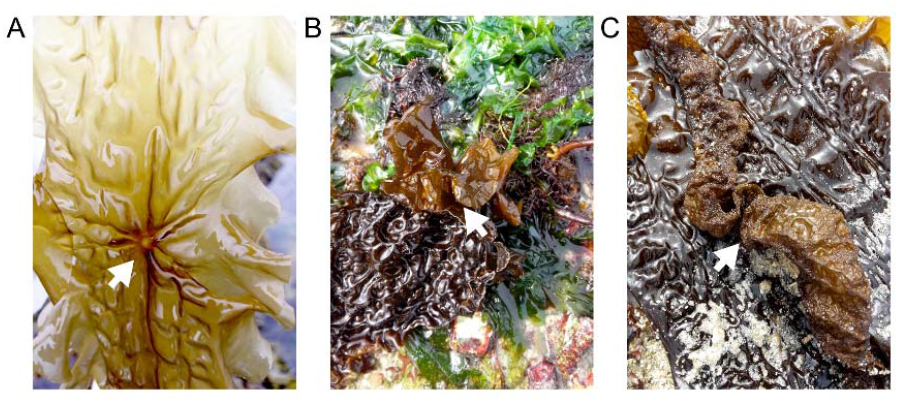
Examples of symptoms observed on “diseased” *S. latissima* individuals. A) hole, B) twisted blade, and C) bubbling in blade

**Figure 6.**
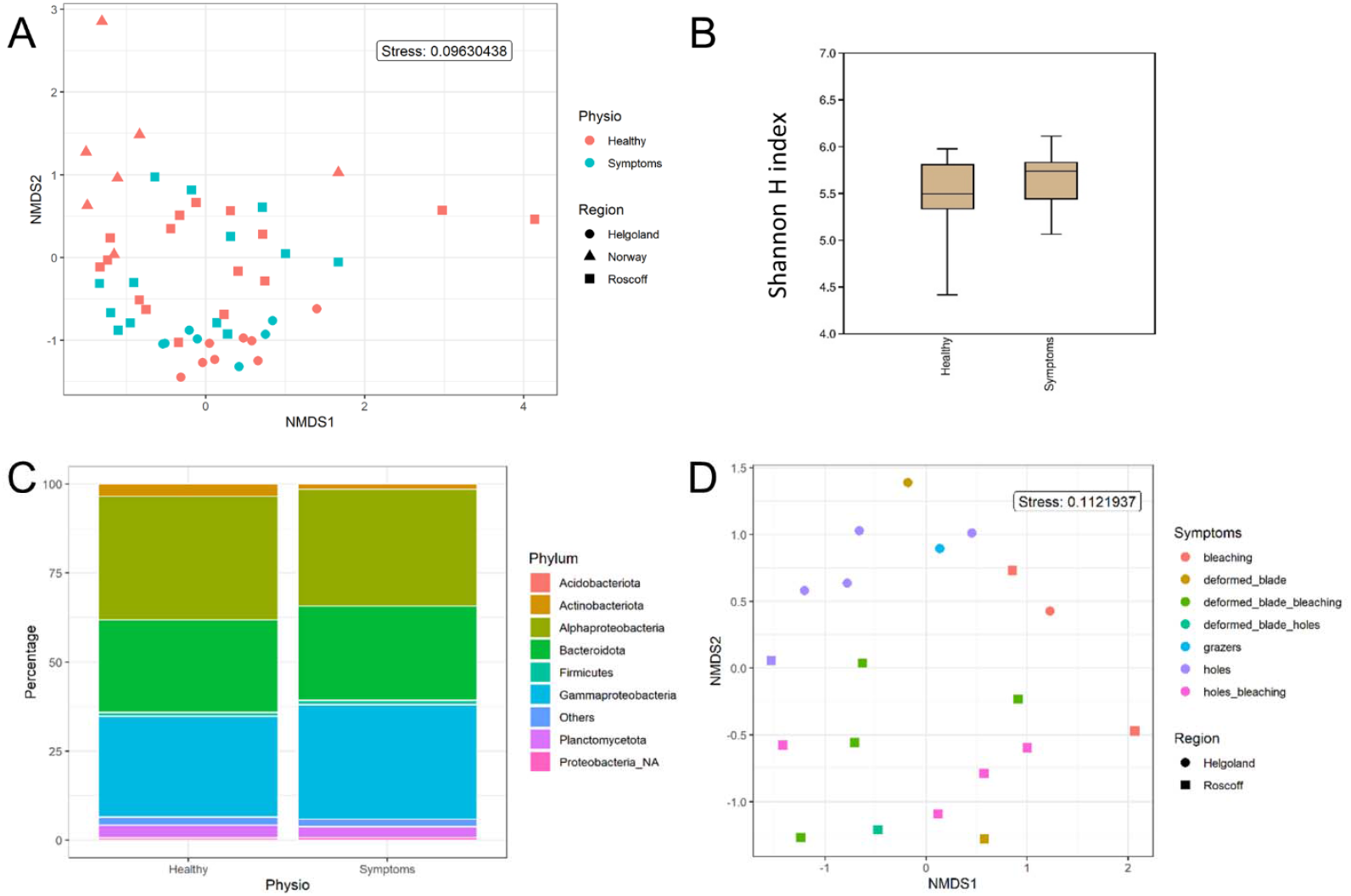
Analysis of samples with symptoms. A) NMDS analysis of the microbiome composition. Results do not show a separation of the healthy and symptomatic samples. B) Box plot of alpha-diversity (Shannon H index) across different sample types. *p-value*=*0.046*. C) Microbiome composition of healthy and symptoms samples. Distribution of 16S rRNA gene metabarcoding sequences per phylum. *Proteobacteria_NA*: not classified as Alpha-or Gammaproteobacteria. D) NMDS analysis of the microbiome composition depending on the symptoms.

## IV. DISCUSSION

Understanding holobiont functioning requires knowledge of the holobiont’s bacterial component. Here we studied the diversity and composition of bacterial communities of *S. latissima* by 16S metabarcoding analysis. The impacts of several factors on bacterial communities were examined: algal blade part, origin of the host, season, and host condition.

### The blade part is the primary driver of samples separation

Distinct bacterial communities were demonstrated to be associated with different parts of *S. latissima* by 16S metabarcoding, and this is the primary factor of separation regardless of region, season, and physiology. Staufenberger et al. (2008) found the same dynamics when working on several *S. latissima* tissue from the Baltic Sea and the North Sea, sampled in winter and spring.

*S. latissima’s* type of growth can explain this difference in bacterial communities. *S. latissima* is a short-lived but perennial species, and growth occurs mainly in the meristem region. From there, the proliferating cells form the thallus. Young algae only have short blades and no access to the surrounding sediment as they only stand upright in the water column. As the thallus grows, it becomes heavier, and the apex finally bends and touches the ground (Kain, 1979; Lüning, 1991). Water currents move the old blade, resulting in access to nearby substrates and a broader environment. Mechanical stress also occurs in this part, which becomes vulnerable to bacterial decomposition, offering new ecological niches for different bacteria (Bengtsson and Øvreås, 2010). Therefore, the younger meristem tissues are typically less colonised by bacteria and exhibit lower bacterial diversity, as previously found (Staufenberger et al., 2008; Goecke et al., 2010; Ihua et al., 2020; Lemay et al., 2021). Furthermore, the synthesis or release of compounds that either have an antimicrobial effect or act as nutrients for the bacteria may vary between the different parts of the blade. This was described for phenolic substances in the kelp *L. hyperborea* and likely contributed to differences in the microbial composition (Bengtsson et al., 2012).

We found a higher proportion of *Planctomycetes* at the apex and *Actinobacteriota* almost exclusively in the meristem. Both phyla are classically found in brown macroalgae (Hollants et al., 2013) and on *S. latissima* [apex: (Staufenberger et al., 2008); meristem: (Tourneroche et al., 2020)]. *Planctomycetes* contain many sulfatase genes (Wegner et al., 2013), which help degrade sulphated polysaccharides. They may also be involved in degrading polysaccharides from the extracellular matrix of microbial biofilms (Parrot et al., 2019), which may explain their higher relative abundance in older tissues that exhibit first signs of degradation. *Actinobacteriota*, on the other hand, is a diverse phylum that has successfully colonised a wide range of habitats (Ul-Hassan and Wellington, 2009), but we currently do not know what features make them successful colonisers of *the S. latissima* meristem.

The bacterial core also reveals the shift from a low to a higher diversity as the blade ages, with four genera found in both algal parts and five additional genera found in >90% of apex samples. Those taxa were also found on the meristem part of *L. digitata* (Ihua et al., 2020) and on the blade of *L. setchellii* (Lemay et al., 2021), *Taonia atomaria* (Paix et al., 2021), and *U. lactuca* (Comba González et al., 2021). Our results are furthermore consistent with the core microbiota found in *S. latissima* and *L. hyperborea* in the United Kingdom (King et al., 2022). *Granulosicoccus* sp. (*Gammaproteobacteria)* was one of the most abundant genera (12% of total reads) and included several ASVs overexpressed in the meristem samples. This genus was also found abundantly on the youngest parts of the sister species *S. japonica* (Balakirev et al., 2012; Zhang et al., 2020) and other kelps like *L. setchellii* (Lemay et al., 2021), *L. hyperborea* (Bengtsson et al., 2012), *Macrocystis pyrifera*, and *Nereocystis luetkeana* (Weigel and Pfister, 2019; Ramírez-Puebla et al., 2022), reinforcing the idea of a strong association between *Granulosicoccus* and the kelp tissue. In the same vein, the genus *Algitalea (Flavobacteriaceae;* 3% of total reads) was one of the “apex” bacterial core genera. This genus belonged to the pioneer bacterial communities found on the apical parts of *T. atomaria* (Paix et al., 2020). Also, the *Flavobacterium* lineage has been recognised as necessary in the decomposition processes of organic matter during algae blooms (Riemann et al., 2000; Pinhassi et al., 2004) and thus may participate in the decay process occurring at the algal apices.

### Regional specificities: tides, seawater, and genetic background

Aside from the apex/meristem duality, the region of origin also was a decisive separation factor. Lachnit et al. (2009) have already shown this region-dependent separation by comparing global epibacterial communities of algae from the North and Baltic Seas. In their study, only *S. latissima* showed regional differences within conspecific algae (contrary to the two other studied *Phaeophyceae*). At a larger geographical scale, results on *Ulva* sp. and *Agarophyton vermiculosum* (Roth-Schulze et al., 2018; Bonthond et al., 2020) suggested that the seaweed microbiota composition, diversity, and functions strongly depend on the local scale, but also shows that processes are acting at larger scales to shape this microbial community, and they need to be identified.

In our study, the regional differences were more pronounced than the seasonal differences obtained for one sampling site, suggesting that region and not just variability between sampling dates was the driving factor. The regional differences might be due to several abiotic factors. The tidal ranges for the three regions, for instance, decrease going north: 10 meters in Roscoff, 3 meters in Helgoland, and less than 1 meter in Skagerrak. In the same vein, winter seawater temperatures are lower in the north, which might favour psychrophilic strains over mesophilic communities, as found in a culture-based study on the surface bacteria of *L. longicruris* (Laycock, 1974). Also, increasing time exposure to rain, wind, or sunlight (UV), waves, currents, hydrostatic pressure, pH, and salinity can lead to cellular stress and, by extension, senescence. This changing environment can offer new ecological niches for different bacteria. Lastly, bacterial communities can be altered by nutrient supply, interspecies competition, and viral infection (Fuhrman et al., 2015; Stal and Cretoiu, 2016).

However, the regional differences might also be due, in part, to the genetic diversity of the algae. Using single nucleotide polymorphisms (SNPs) and microsatellites, Guzinski et al. (2016, 2020) determined that *S. latissima* individuals from Roscoff, Helgoland, and Norway are genetically distinct, and this might lead to the attraction of different bacterial species (Griffiths et al., 2019). Furthermore, *S. latissima* displays a unique lipidomic signature depending on its geographic origin (Monteiro et al., 2020), as the content of chemical elements (C, H, N, S), fatty acids, and lipids varies depending on the region. These molecules are common components of membranes (Harwood, 2004) and might influence the attractiveness of the algal surface for several bacterial strains.

### Shifts in bacterial communities depending on the season

The third factor we examined was seasonality. Laycock (1974) observed that as the seasonal temperatures decreased, the bacterial communities of *L. longicruris* shifted from mesophilic to psychrophilic strains. When working with *L. hyperborea*, Bengtsson et al. (2010) hypothesised that the seasonal succession in the bacterial communities might be explained by abiotic factors like seawater temperature and biotic factors like seasonal changes in the kelp substrate. Indeed, seawater temperature alone does not seem to have been the most important factor in our data, as the seawater was coldest in winter and spring (<12°C), but the samples from autumn and winter were more similar in their bacterial communities. Other physicochemical parameters might also play a role. Nitrogen is an important element for organisms, and microorganisms can take up nitrogen in different forms such as nitrate, nitrite, ammonium, urea, organic nitrogen, and in some cases, dinitrogen gas (N2), depending on the organism (Zehr and Ward, 2002). Nitrate, nitrite, and ammonium concentrations follow seasonal variations, with nitrates being lower in summer and higher in winter and nitrites at their highest in autumn. Phosphorus is another essential nutrient for primary production in the euphotic zone. Most of the phosphorus is present in the oxidised form as free phosphate or bound to organic matter. The phosphate concentration was lowest in springtime. Several ASVs overrepresented in spring belong to the *Roseobacter* clade, and Atlantic strains of this genus are known to possess abundant high-affinity phosphorus uptake systems, constituting likely adaptations to low environmental phosphate concentrations (Newton et al., 2010).

Lastly, seasonal variations may also be due to seasonal changes in the alga’s chemical composition. For instance, Schiener et al. (2015) demonstrated that in *S. latissima*, polyphenol levels are higher between May and July and then decrease, reaching their lowest in March. This could be interpreted as a defence against bacterial colonisation as the seawater temperature rises, polyphenols being known for their wide range of antimicrobial properties (Zhang et al., 2006; Daglia, 2012). Also, carbohydrate content (laminarin and mannitol) is higher in summer (Schiener et al., 2015), and these are both substrates easy to degrade by the bacteria (Alderkamp et al., 2007; Jeske et al., 2013; Groisillier et al., 2015). Similarly, algal iodine content is generally lower in summer (Nitschke et al., 2018), and the algae’s production of toxic iodine compounds may control the surface biofilm and repulse microbial pathogens (Rodeheaver et al., 1982; Gobet et al., 2017).

### Is there a microbial signature characteristic for algae in poor health?

Some bacteria affect the alga in a deleterious manner by decomposing cell material, like alginate and laminarin (Laycock, 1974; Dimitrieva and Dimitriev, 1997; Sawabe et al., 1998b; Ivanova et al., 2003) or by causing diseases like *Alteromonas* species (Vairappan et al., 2001; Peng and Li, 2013) and species of *Pseudoalteromonas* (Sawabe et al., 1998a). Ihua et al. (2019) have shown that the microbial communities (phyla level) associated with intact *Ascophyllum* differ from rotting algae, suggesting that the decay process might shape the associated bacterial community. Similarly, the microbial communities of *Ecklonia* are strongly associated with the algal condition (stressed or not) more than with other variables (Marzinelli et al., 2015). Moreover, the core bacterial community characteristic of healthy algae may be lost when hosts are subjected to stress, and the microbiota of stressed individuals of *Ecklonia* were more similar to each other at a given location than those on healthy hosts (Marzinelli et al., 2015); which is contrary to the so-called Anna Karenina principle (Zaneveld et al., 2017; Ma, 2020), stating that all “healthy” microbiomes are alike and each “symptom” microbiome is “sick” in its own way.

These data reinforce our hypothesis of a characteristic microbial signature for algae in poor health, regardless of the specific symptoms. In our study, the changes in bacterial communities between healthy and diseased individuals are visible only at a lower taxonomic scale, and we found 9 ASVs that were differentially expressed between the healthy (5 ASVs) and diseased samples (4 ASVs). ASVs characteristic for the latter belong to the genus *Tenacibaculum* and the *Alteromonadales*, known for their alginate lyase activities (Thomas et al., 2021), and to the *Roseobacter* clade, known for their production of quorum-sensing molecules, a phenomenon involved in virulence and pathogenicity (Buchan et al., 2005; Wagner-Döbler and Biebl, 2006; Brinkhoff et al., 2008). One of the healthy specific ASVs is a *Granulosicoccus* sp., emphasising the importance of this genus in the algal microbiota. The fact that several ASVs were found to differ indicates that, regardless of the type of disease, an alga that is not well will undergo characteristic changes in the microbiome. Moreover, these ASVs signatures are probably stable because they are derived from different places, times of the year, and symptoms, as shown for *Ecklonia* (Marzinelli et al., 2015). Although this would require additional developments, these signatures might also be helpful as bioindicators for kelp health.

## V. CONCLUSION

In conclusion, our study provides an extensive overview of the *S. latissima* microbiome and highlights several factors driving its variability. In particular, the observation that the blade part had a more profound impact on the microbial composition than season or region, both of which are associated with changes in the abiotic environment, underlines the extent to which algal hosts select their associated microbiota. Our discovery of microbial signatures characteristic for diseased *S. latissima* individuals that persist in our dataset independently of the disease symptoms further supports this hypothesis. Given the variety of symptoms observed in our samples, it is unlikely that the same bacteria could be the causative agents in all cases. Rather, the different types of disease likely cause similar changes in the host, which would lead to similar microbial changes. Understanding these signatures will be of interest for fundamental research on the different algal diseases; in the long run, it may also help develop molecular markers of host health to survey natural populations or aquacultures.

## DECLARATIONS

### Data availability

Raw sequence data were deposited at the European Nucleotide Archive under project accession number ENA: PRJEB47035.

### Code availability

not applicable

### Authors’ contributions

Designed study: BBD, SD; Sampling: BBD, SF, SD; Performed experiments: BBD, EL, GT; Analysed data: BBD, SD; Wrote the manuscript: BBD, SD; Provided valuable input and corrected the manuscript: CB.

## ACKNOWLEDGEMENTS

We thank François Thomas, Maéva Brunet, and Nolwen Le Duff for their primers and for providing the mock community; Sylvie Rousvoal for advice on sample preparation, Jonas Collén for advice and participating in the first sampling, Kai Bishof and Nora Diehl for providing samples from Svalbard, which unfortunately could not be included, and Catherine Leblanc for helpful discussions. This work benefited from access to the Station Biologique de Roscoff, an EMBRC-France and EMBRC-ERIC Site.

## SUPPLEMENTARY DATA

**Table S1 - Taxonomic affiliations of over-expressed ASVs for each comparison** (algal part, regions, seasons and symptoms)

**Figure S1.**
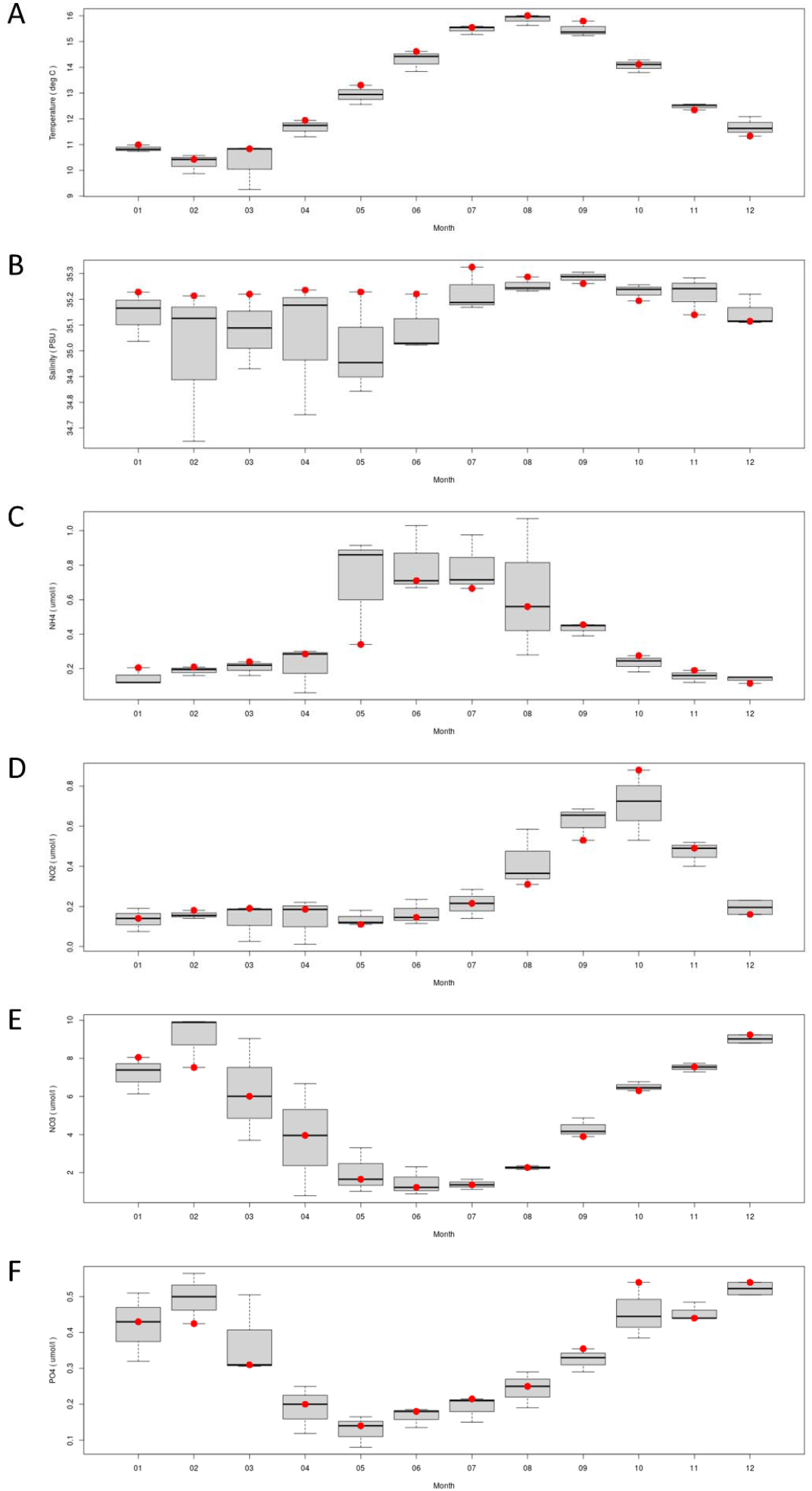
Seasonal variations in A) temperature, B) salinity and C) ammonium, D) nitrites, E) nitrates, and F) phosphate concentrations. Roscoff, 2019. Legend for each month, rectangle: region inside 1^st^ and 3^rd^ quartiles, bold line: median value, dashed error bars: 1^st^ and 9^th^ deciles, red point: value for the selected year.

## REFERENCES

Alderkamp, A.-C., Van Rijssel, M., and Bolhuis, H. (2007). Characterization of marine bacteria and the activity of their enzyme systems involved in degradation of the algal storage glucan laminarin: FEMS Microbiol. Ecol. 59, 108–117. doi: 10.1111/j.1574-6941.2006.00219.x.

Araújo, R. M., Assis, J., Aguillar, R., Airoldi, L., Bárbara, I., Bartsch, I., et al. (2016). Status, trends and drivers of kelp forests in Europe: an expert assessment. Biodivers. Conserv. 25, 1319–1348. doi: 10.1007/s10531-016-1141-7.

Balakirev, E. S., Krupnova, T. N., and Ayala, F. J. (2012). Symbiotic associations in the phenotypically-diverse brown alga *Saccharina japonica*. PLoS ONE 7, e39587. doi: 10.1371/journal.pone.0039587.

Bengtsson, M. M., and Øvreås, L. (2010). *Planctomycetes* dominate biofilms on surfaces of the kelp *Laminaria hyperborea*. BMC Microbiol. 10, 261. doi: 10.1186/1471-2180-10-261.

Bengtsson, M. M., Sjøtun, K., Lanzén, A., and Øvreås, L. (2012). Bacterial diversity in relation to secondary production and succession on surfaces of the kelp *Laminaria hyperborea*. ISME J. 6, 2188–2198. doi: 10.1038/ismej.2012.67.

Bengtsson, M., Sjøtun, K., and Øvreås, L. (2010). Seasonal dynamics of bacterial biofilms on the kelp *Laminaria hyperborea*. Aquat. Microb. Ecol. 60, 71–83. doi: 10.3354/ame01409.

Benjamini, Y., and Hochberg, Y. (1995). Controlling the false discovery rate: a practical and powerful approach to multiple testing. J. R. Stat. Soc. Ser. B Methodol. 57, 289–300. doi: 10.1111/j.2517-6161.1995.tb02031.x.

Bernard, M., Rousvoal, S., Jacquemin, B., Ballenghien, M., Peters, A. F., and Leblanc, C. (2017). qPCR-based relative quantification of the brown algal endophyte *Laminarionema elsbetiae* in *Saccharina latissima*: variation and dynamics of host - endophyte interactions. J. Appl. Phycol. 30, 2901–2911. doi: 10.1007/s10811-017-1367-0.

Bolton, J. J. (2010). The biogeography of kelps (*Laminariales, Phaeophyceae*): a global analysis with new insights from recent advances in molecular phylogenetics. Helgol. Mar. Res. 64, 263–279. doi: 10.1007/s10152-010-0211-6.

Bonthond, G., Bayer, T., Krueger-Hadfield, S. A., Barboza, F. R., Nakaoka, M., Valero, M., et al. (2020). How do microbiota associated with an invasive seaweed vary across scales? Mol. Ecol. 29, 2094–2108. doi:10.1111/mec.15470.

Bosch, T. C. G., and Miller, D. J. (2016). “Bleaching as an obvious dysbiosis in corals,” in The Holobiont Imperative (Vienna: Springer Vienna), 113–125. doi: 10.1007/978-3-7091-1896-2_9.

Bourne, D., Iida, Y., Uthicke, S., and Smith-Keune, C. (2008). Changes in coral-associated microbial communities during a bleaching event. ISME J. 2, 350–363. doi: 10.1038/ismej.2007.112.

Brinkhoff, T., Giebel, H.-A., and Simon, M. (2008). Diversity, ecology, and genomics of the Roseobacter clade: a short overview. Arch. Microbiol. 189, 531–539. doi: 10.1007/s00203-008-0353-y.

Buchan, A., González, J. M., and Moran, M. A. (2005). Overview of the Marine Roseobacter Lineage. Appl. Environ. Microbiol. 71, 5665–5677. doi: 10.1128/AEM.71.10.5665-5677.2005.

Burgunter-Delamare, B., KleinJan, H., Frioux, C., Fremy, E., Wagner, M., Corre, E., et al. (2020). Metabolic complementarity between a brown alga and associated cultivable bacteria provide indications of beneficial interactions. Front. Mar. Sci. 7, 85. doi: 10.3389/fmars.2020.00085.

Callahan, B. J., McMurdie, P. J., Rosen, M. J., Han, A. W., Johnson, A. J. A., and Holmes, S. P. (2016). DADA2: High-resolution sample inference from Illumina amplicon data. Nat. Methods 13, 581–583. doi: 10.1038/nmeth.3869.

Comba González, N. B., Niño Corredor, A. N., López Kleine, L., and Montoya Castaño, D. (2021). Temporal changes of the epiphytic bacteria community from the marine macroalga *Ulva lactuca* (Santa Marta, Colombian-Caribbean). Curr. Microbiol. 78, 534–543. doi: 10.1007/s00284-020-02302-x.

Corre, S., and Prieur, D. (1990). Density and morphology of epiphytic bacteria on the kelp *Laminaria digitata*. Bot. Mar. 33. doi: 10.1515/botm.1990.33.6.515.

Daglia, M. (2012). Polyphenols as antimicrobial agents. Curr. Opin. Biotechnol. 23, 174–181. doi: 10.1016/j.copbio.2011.08.007.

Dimitrieva, G. Y., and Dimitriev, S. M. (1997). Symbiotic microflora of brown algae of the genus *Laminaria* as a bioindicator of the ecological condition of coastal laminarian biocenoses. Oceanogr. Lit. Rev. 11, 1330.

Dittami, S. M., Duboscq-Bidot, L., Perennou, M., Gobet, A., Corre, E., Boyen, C., et al. (2016). Host-microbe interactions as a driver of acclimation to salinity gradients in brown algal cultures. ISME J. 10, 51–63. doi: 10.1038/ismej.2015.104.

Egan, S., Harder, T., Burke, C., Steinberg, P., Kjelleberg, S., and Thomas, T. (2013). The seaweed holobiont: understanding seaweed–bacteria interactions. FEMS Microbiol. Rev. 37, 462–476. doi: 10.1111/1574-6976.12011.

Fuhrman, J. A., Cram, J. A., and Needham, D. M. (2015). Marine microbial community dynamics and their ecological interpretation. Nat. Rev. Microbiol. 13, 133–146. doi: 10.1038/nrmicro3417.

Gobet, A., Corre, E., Correc, G., Delage, L., Dittami, S., KleinJan, H., et al. (2017). Characterization of the epiphytic bacterial community associated with the kelp *Laminaria digitate*. Phycologia 56, 64.

Goecke, F., Labes, A., Wiese, J., and Imhoff, J. (2010). Chemical interactions between marine macroalgae and bacteria. Mar. Ecol. Prog. Ser. 409, 267–299. doi: 10.3354/meps08607.

Griffiths, S. M., Antwis, R. E., Lenzi, L., Lucaci, A., Behringer, D. C., Butler, M. J., et al. (2019). Host genetics and geography influence microbiome composition in the sponge *Ircinia campana*. J. Anim. Ecol. 88, 1684–1695. doi: 10.1111/1365-2656.13065.

Groisillier, A., Labourel, A., Michel, G., and Tonon, T. (2015). The mannitol utilization system of the marine bacterium *Zobellia galactanivorans*. Appl. Environ. Microbiol. 81, 1799–1812. doi: 10.1128/AEM.02808-14.

Guzinski, J., Mauger, S., Cock, J. M., and Valero, M. (2016). Characterization of newly developed expressed sequence tag-derived microsatellite markers revealed low genetic diversity within and low connectivity between European *Saccharina latissima* populations. J. Appl. Phycol. 28, 3057–3070. doi: 10.1007/s10811-016-0806-7.

Guzinski, J., Ruggeri, P., Ballenghien, M., Mauger, S., Jacquemin, B., Jollivet, C., et al. (2020). Seascape genomics of the sugar kelp *Saccharina latissima* along the North Eastern Atlantic Latitudinal Gradient. Genes 11, 1503. doi: 10.3390/genes11121503.

Hammer, Ø., Harper, D. A., and Ryan, P. D. (2001). PAST: Paleontological statistics software package for education and data analysis. Palaeontol. Electron. 4, 9.

Harwood, J. L. (2004). “Membrane lipids in algae,” in Lipids in Photosynthesis: Structure, Function and Genetics Advances in Photosynthesis and Respiration., eds. S. Paul-André and M. Norio (Dordrecht: Kluwer Academic Publishers), 53–64. doi: 10.1007/0-306-48087-5_3.

Hollants, J., Leliaert, F., De Clerck, O., and Willems, A. (2013). What we can learn from sushi: a review on seaweed–bacterial associations. FEMS Microbiol. Ecol. 83, 1–16. doi: 10.1111/j.1574-6941.2012.01446.x.

Ihua, M., Guihéneuf, F., Mohammed, H., Margassery, L., Jackson, S., Stengel, D., et al. (2019). Microbial population changes in decaying *Ascophyllum nodosum* result in macroalgal-polysaccharide-degrading bacteria with potential applicability in enzyme-assisted extraction technologies. Mar. Drugs 17, 200. doi: 10.3390/md17040200.

Ihua, M. W., FitzGerald, J. A., Guihéneuf, F., Jackson, S. A., Claesson, M. J., Stengel, D. B., et al. (2020). Diversity of bacteria populations associated with different thallus regions of the brown alga *Laminaria digitata*. PLOS ONE 15, e0242675. doi: 10.1371/journal.pone.0242675.

Illumina (2013). 16S Metagenomic sequencing library preparation. Prep. 16S Ribosomal RNA Gene Amplicons Illumina MiSeq Syst., 1–28.

Ivanova, E. P., Bakunina, I. Yu., Nedashkovskaya, O. I., Gorshkova, N. M., Alexeeva, Y. V., Zelepuga, E. A., et al. (2003). Ecophysiological variabilities in ectohydrolytic enzyme activities of some *Pseudoalteromonas* species, *P. citrea, P. issachenkonii*, and *P. nigrifaciens*. Curr. Microbiol. 46, 6–10. doi: 10.1007/s00284-002-3794-6.

Jeske, O., Jogler, M., Petersen, J., Sikorski, J., and Jogler, C. (2013). From genome mining to phenotypic microarrays: *Planctomycetes* as source for novel bioactive molecules. Antonie Van Leeuwenhoek 104, 551–567. doi: 10.1007/s10482-013-0007-1.

Kain, J. M. (1979). A view of the genus *Laminaria*. Oceanogr. Mar. Biol. Annu. Rev. 17, 101–161.

King, N. G., Moore, P. J., Thorpe, J. M., and Smale, D. A. (2022). Consistency and Variation in the Kelp Microbiota: Patterns of Bacterial Community Structure Across Spatial Scales. Microb. Ecol. doi: 10.1007/s00248-022-02038-0.

Lachnit, T., Blümel, M., Imhoff, J., and Wahl, M. (2009). Specific epibacterial communities on macroalgae: phylogeny matters more than habitat. Aquat. Biol. 5, 181–186. doi: 10.3354/ab00149.

Laycock, R. A. (1974). The detrital food chain based on seaweeds. I. Bacteria associated with the surface of *Laminaria* fronds. Mar. Biol. 25, 223–231. doi: 10.1007/BF00394968.

Lemay, M. A., Davis, K. M., Martone, P. T., and Parfrey, L. W. (2021). Kelp-associated microbiota are structured by host anatomy. J. Phycol. 57, 1119–1130. doi: 10.1111/jpy.13169.

Lin, H., and Peddada, S. D. (2020). Analysis of compositions of microbiomes with bias correction. Nat. Commun. 11, 3514. doi: 10.1038/s41467-020-17041-7.

Lüning, K. (1991). Seaweeds: their environment, biogeography, and ecophysiology. New York, NY: Wiley.

Ma, Z. (Sam) (2020). Testing the Anna Karenina Principle in Human Microbiome-Associated Diseases. iScience 23, 101007. doi: 10.1016/j.isci.2020.101007.

Marzinelli, E. M., Campbell, A. H., Zozaya Valdes, E., Vergés, A., Nielsen, S., Wernberg, T., et al. (2015). Continental-scale variation in seaweed host-associated bacterial communities is a function of host condition, not geography. Environ. Microbiol. 17, 4078–4088. doi: 10.1111/1462-2920.12972.

Mazure, H. G. F., and Field, J. G. (1980). Density and ecological importance of bacteria on kelp fronds in an upwelling region. J. Exp. Mar. Biol. Ecol. 43, 173–182. doi: 10.1016/0022-0981(80)90024-6.

McHugh, D. J. (2003). A guide to the seaweed industry. Rome: Food and Agriculture Organization of the United Nations Available at: https://www.fao.org/3/y4765e/y4765e.pdf.

McMurdie, P. J., and Holmes, S. (2013). phyloseq: an R package for reproducible interactive analysis and graphics of microbiome census data. PLoS ONE 8, e61217. doi: 10.1371/journal.pone.0061217.

Monteiro, J. P., Rey, F., Melo, T., Moreira, A. S. P., Arbona, J.-F., Skjermo, J., et al. (2020). The unique lipidomic signatures of *Saccharina latissima* can be used to pinpoint their geographic origin. Biomolecules 10, 107. doi: 10.3390/biom10010107.

Newton, R. J., Griffin, L. E., Bowles, K. M., Meile, C., Gifford, S., Givens, C. E., et al. (2010). Genome characteristics of a generalist marine bacterial lineage. ISME J. 4, 784–798. doi: 10.1038/ismej.2009.150.

Nitschke, U., Walsh, P., McDaid, J., and Stengel, D. B. (2018). Variability in iodine in temperate seaweeds and iodine accumulation kinetics of *Fucus vesiculosus* and *Laminaria digitata* (Phaeophyceae, Ochrophyta). J. Phycol. 54, 114–125. doi: 10.1111/jpy.12606.

Paix, B., Carriot, N., Barry-Martinet, R., Greff, S., Misson, B., Briand, J.-F., et al. (2020). A multi-omics analysis suggests links between the differentiated surface metabolome and epiphytic microbiota along the thallus of a Mediterranean seaweed holobiont. Front. Microbiol. 11, 494. doi: 10.3389/fmicb.2020.00494.

Paix, B., Layglon, N., Le Poupon, C., D’Onofrio, S., Misson, B., Garnier, C., et al. (2021). Integration of spatio-temporal variations of surface metabolomes and epibacterial communities highlights the importance of copper stress as a major factor shaping host-microbiota interactions within a Mediterranean seaweed holobiont. Microbiome 9, 201. doi: 10.1186/s40168-021-01124-8.

Parrot, D., Blümel, M., Utermann, C., Chianese, G., Krause, S., Kovalev, A., et al. (2019). Mapping the surface microbiome and metabolome of brown seaweed *Fucus vesiculosus* by amplicon sequencing, integrated metabolomics and imaging techniques. Sci. Rep. 9, 1061. doi: 10.1038/s41598-018-37914-8.

Peixoto, R. S., Rosado, P. M., Leite, D. C. de A., Rosado, A. S., and Bourne, D. G. (2017). Beneficial Microorganisms for Corals (BMC): Proposed Mechanisms for Coral Health and Resilience. Front. Microbiol. 8. doi: 10.3389/fmicb.2017.00341.

Peng, Y., and Li, W. (2013). A bacterial pathogen infecting gametophytes of *Saccharina japonica (Laminariales, Phaeophyceae)*. Chin. J. Oceanol. Limnol. 31, 366–373. doi: 10.1007/s00343-013-2136-9.

Peteiro, C. (2018). “Alginate production from marine macroalgae, with emphasis on kelp farming,” in Alginates and their biomedical applications Springer Series in Biomaterials Science and Engineering., eds. B. H. A. Rehm and M. F. Moradali (Singapore: Springer Singapore), 27–66. doi: 10.1007/978-981-10-6910-9_2.

Pinhassi, J., Sala, M. M., Havskum, H., Peters, F., Guadayol, Ò., Malits, A., et al. (2004). Changes in bacterioplankton composition under different phytoplankton regimens. Appl. Environ. Microbiol. 70, 6753–6766. doi: 10.1128/AEM.70.11.6753-6766.2004.

Ramírez-Puebla, S. T., Weigel, B. L., Jack, L., Schlundt, C., Pfister, C. A., and Mark Welch, J. L. (2022). Spatial organization of the kelp microbiome at micron scales. Microbiome 10, 52. doi: 10.1186/s40168-022-01235-w.

Riemann, L., Steward, G. F., and Azam, F. (2000). Dynamics of bacterial community composition and activity during a mesocosm diatom bloom. Appl. Environ. Microbiol. 66, 578–587. doi: 10.1128/AEM.66.2.578-587.2000.

Rodeheaver, G., Bellamy, W., Kody, M., Spatafora, G., Fitton, L., Leyden, K., et al. (1982). Bactericidal activity and toxicity of iodine-containing solutions in wounds. Arch. Surg. 117, 181–186. doi: 10.1001/archsurg.1982.01380260051009.

Roth-Schulze, A. J., Pintado, J., Zozaya-Valdés, E., Cremades, J., Ruiz, P., Kjelleberg, S., et al. (2018). Functional biogeography and host specificity of bacterial communities associated with the Marine Green Alga *Ulva* spp. Mol. Ecol.27, 1952–1965. doi: 10.1111/mec.14529.

Roussel, S., Caralp, C., Leblanc, C., Le Grand, F., Stiger-Pouvreau, V., Coulombet, C., et al. (2019). Impact of nine macroalgal diets on growth and initial reproductive investment in juvenile abalone *Haliotis tuberculata*. Aquaculture 513, 734385. doi: 10.1016/j.aquaculture.2019.734385.

Sawabe, T., Makino, H., Tatsumi, M., Nakano, K., Tajima, K., Iqbal, M. M., et al. (1998a). Pseudoalteromonas bacteriolytica sp. nov., a marine bacterium that is the causative agent of red spot disease of *Laminaria japonica*. Int. J. Syst. Bacteriol. 48, 769–774. doi: 10.1099/00207713-48-3-769.

Sawabe, T., Sawada, C., Suzuki, E., and Ezura, Y. (1998b). Intracellular alginate-oligosaccharide degrading enzyme activity that is incapable of degrading intact sodium alginate from a marine bacterium *Alteromonas* sp. Fish. Sci. 64, 320–324. doi: 10.2331/fishsci.64.320.

Schiel, D., and Lilley, S. (2007). Gradients of disturbance to an algal canopy and the modification of an intertidal community. Mar. Ecol. Prog. Ser. 339, 1–11. doi: 10.3354/meps339001.

Schiel, D. R., and Foster, M. S. (2006). The population biology of large brown seaweeds: ecological consequences of multiphase life histories in dynamic coastal environments. Annu. Rev. Ecol. Evol. Syst. 37, 343–372. doi: 10.1146/annurev.ecolsys.37.091305.110251.

Schiener, P., Black, K. D., Stanley, M. S., and Green, D. H. (2015). The seasonal variation in the chemical composition of the kelp species *Laminaria digitata, Laminaria hyperborea, Saccharina latissima* and *Alaria esculenta*. J. Appl. Phycol. 27, 363–373. doi: 10.1007/s10811-014-0327-1.

Smale, D. A. (2020). Impacts of ocean warming on kelp forest ecosystems. New Phytol. 225, 1447–1454. doi: 10.1111/nph.16107.

Smit, A. J. (2004). Medicinal and pharmaceutical uses of seaweed natural products: A review. J. Appl. Phycol. 16, 245–262. doi: 10.1023/B:JAPH.0000047783.36600.ef.

Stal, L. J., and Cretoiu, M. S. eds. (2016). The Marine Microbiome. Cham: Springer International Publishing doi: 10.1007/978-3-319-33000-6.

Staufenberger, T., Thiel, V., Wiese, J., and Imhoff, J. F. (2008). Phylogenetic analysis of bacteria associated with *Laminaria saccharina*. FEMS Microbiol. Ecol. 64, 65–77. doi: 10.1111/j.1574-6941.2008.00445.x.

Thomas, F., Dittami, S. M., Brunet, M., Le Duff, N., Tanguy, G., Leblanc, C., et al. (2019). Evaluation of a new primer combination to minimize plastid contamination in 16S rDNA metabarcoding analyses of alga-associated bacterial communities. Environ. Microbiol. Rep. 12, 30–37. doi: 10.1111/1758-2229.12806.

Tourneroche, A., Lami, R., Burgaud, G., Domart-Coulon, I., Li, W., Gachon, C., et al. (2020). The bacterial and fungal microbiota of *Saccharina latissima* (Laminariales, Phaeophyceae). Front. Mar. Sci. 7, 587566. doi: 10.3389/fmars.2020.587566.

Ul-Hassan, A., and Wellington, E. M. (2009). “Actinobacteria,” in Encyclopedia of Microbiology (Elsevier), 25–44. doi: 10.1016/B978-012373944-5.00044-4.

Vairappan, C. S., Suzuki, M., Motomura, T., and Ichimura, T. (2001). Pathogenic bacteria associated with lesions and thallus bleaching symptoms in the Japanese kelp *Laminaria religiosa* Miyabe (Laminariales, Phaeophyceae). Hydrobiologia 445, 183–191. doi: 10.1023/A:1017517832302.

Wagner-Döbler, I., and Biebl, H. (2006). Environmental Biology of the Marine *Roseobacter* Lineage. Annu. Rev. Microbiol. 60, 255–280. doi: 10.1146/annurev.micro.60.080805.142115.

Wegner, C.-E., Richter-Heitmann, T., Klindworth, A., Klockow, C., Richter, M., Achstetter, T., et al. (2013). Expression of sulfatases in *Rhodopirellula baltica* and the diversity of sulfatases in the genus *Rhodopirellula*. Mar. Genomics 9, 51–61. doi: 10.1016/j.margen.2012.12.001.

Weigel, B. L., and Pfister, C. A. (2019). Successional dynamics and seascape-level patterns of microbial communities on the canopy-forming kelps *Nereocystis luetkeana* and *Macrocystis pyrifera*. Front. Microbiol. 10, 346. doi: 10.3389/fmicb.2019.00346.

Wiese, J., Thiel, V., Nagel, K., Staufenberger, T., and Imhoff, J. F. (2009). Diversity of antibiotic-active bacteria associated with the brown alga *Laminaria saccharina* from the Baltic Sea. Mar. Biotechnol. 11, 287–300. doi: 10.1007/s10126-008-9143-4.

Zaneveld, J. R., McMinds, R., and Vega Thurber, R. (2017). Stress and stability: applying the Anna Karenina principle to animal microbiomes. Nat. Microbiol. 2, 17121. doi: 10.1038/nmicrobiol.2017.121.

Zehr, J. P., and Ward, B. B. (2002). Nitrogen Cycling in the Ocean: New Perspectives on Processes and Paradigms. Appl. Environ. Microbiol. 68, 1015–1024. doi: 10.1128/AEM.68.3.1015-1024.2002.

Zhang, Q., Zhang, J., Shen, J., Silva, A., Dennis, D. A., and Barrow, C. J. (2006). A simple 96-well microplate method for estimation of total polyphenol content in seaweeds. J. Appl. Phycol. 18, 445–450. doi: 10.1007/s10811-006-9048-4.

Zhang, R., Chang, L., Xiao, L., Zhang, X., Han, Q., Li, N., et al. (2020). Diversity of the epiphytic bacterial communities associated with commercially cultivated healthy and diseased *Saccharina japonica* during the harvest season. J. Appl. Phycol. 32, 2071–2080. doi: 10.1007/s10811-019-02025-y.

